# Meaning maps and saliency models based on deep convolutional neural networks are insensitive to image meaning when predicting human fixations

**DOI:** 10.1101/840256

**Authors:** Marek A. Pedziwiatr, Matthias Kümmerer, Thomas S.A. Wallis, Matthias Bethge, Christoph Teufel

**Affiliations:** Cardiff University, Cardiff University Brain Research Imaging Centre (CUBRIC), School of Psychology Cardiff, United Kingdom; University of Tübingen, Center for Integrative Neuroscience, Tübingen, Germany; Bernstein Center for Computational Neuroscience, Tübingen, Germany

**Keywords:** eye movements, natural scenes, saliency, deep neural networks, meaning maps

## Abstract

Eye movements are vital for human vision, and it is therefore important to understand how observers decide where to look. Meaning maps (MMs), a technique to capture the distribution of semantic importance across an image, have recently been proposed to support the hypothesis that meaning rather than image features guide human gaze. MMs have the potential to be an important tool far beyond eye-movements research. Here, we examine central assumptions underlying MMs. First, we compared the performance of MMs in predicting fixations to saliency models, showing that DeepGaze II – a deep neural network trained to predict fixations based on high-level features rather than meaning – outperforms MMs. Second, we show that whereas human observers respond to changes in meaning induced by manipulating object-context relationships, MMs and DeepGaze II do not. Together, these findings challenge central assumptions underlying the use of MMs to measure the distribution of meaning in images.

## Introduction

Human eyes resolve fine detail only in a small, central part of the visual field, with resolution dropping off rapidly in the periphery. To sample details, we move our eyes to orient the high-resolution part of our visual system successively to different parts of a visual scene. Information about these small scene parts is extracted during fixations – short periods in which the eyes are relatively stable. Thus, due to the structure of our visual system, human vision depends on eye movements. How the brain decides where to look in a visual scene is therefore an important question. A long-standing hypothesis suggests that semantic content of image regions is important in guiding eye movements. Recent work presented meaning maps (MMs) as a tool to test this hypothesis (Henderson & Hayes, 2017, 2018). This technique aims to index the spatial distribution of meaning across an image, which has potential applications far beyond eye-movement research. Here, we assess and challenge central assumptions of this novel tool.

A classic finding in eye-movement research shows that the specific task of an observer has an influence on where they direct their eyes (Yarbus, 1967; Hayhoe & Ballard, 2005). But in everyday life, we frequently move our eyes without any goal other than to explore the environment. In the lab, this behavior is examined in free-viewing paradigms, during which eye movements are recorded while images are viewed without an explicit task (Koehler, Guo, Zhang, & Eckstein, 2014, but see Tatler, Hayhoe, Land, & Ballard, 2011). To explain what guides eye movements during free viewing, two opposing accounts have been put forward.

According to the first account, eye movements are guided primarily by image characteristics (Borji, Sihite, & Itti, 2013; Itti & Koch, 2001; Parkhurst, Law, & Niebur, 2002). Potential support for this view comes from saliency models: algorithms, which exclusively use visual features of an image to predict human fixations. Although early models, which used only simple features such as local intensity or colors (Itti & Koch, 2000), are now deemed only moderately successful (Bylinskii et al., 2014), more recent saliency models achieve a remarkably high performance (Kümmerer, Wallis, Gatys, & Bethge, 2017). These models harness deep convolutional neural networks – biologically inspired machine learning algorithms, that somewhat resemble the human visual system (Kietzmann, McClure, & Kriegeskorte, 2019). However, even such models rely solely on visual features, albeit high-level ones.

In contrast to the idea underlying saliency models, several authors have argued that during free viewing, eye movements are mainly guided by the semantic content of the visual scene (Henderson, Malcolm, & Schandl, 2009; Nyström & Holmqvist, 2008; Onat, Açik, Schumann, & König, 2014; Rider, Coutrot, Pellicano, Dakin, & Mareschal, 2018; Stoll, Thrun, Nuthmann, & Einhäuser, 2015). This perspective differs fundamentally from the saliency-based approach. Attributing meaning to certain parts of the scene is impossible without prior knowledge of the world, i.e., a factor that is independent of the visual input (Hegde & Kersten, 2010; Teufel, Dakin, & Fletcher, 2018). Consequently, the notion that semantic content guides eye-movements is inconsistent with the idea that the allocation of fixations is dependent solely on the distribution of image features. Given that meaning is not image-computable, the notion that semantic content guides eye-movements is inconsistent with the idea that the eye-movements are dependent solely on the distribution of image features.

A string of recent studies has claimed to provide support for the role of meaning in driving eye movements (Hayes & Henderson, 2019; Henderson & Hayes, 2017, 2018; Henderson, Hayes, Rehrig, & Ferreira, 2018; Peacock, Hayes, & Henderson, 2018). These studies (reviewed in Henderson, Hayes, Peacock, & Rehrig, 2019) are based on a novel technique called meaning maps (MMs). A MM for a given image is created by breaking it down into small isolated patches, which are rated for their meaningfulness independently from the rest of the visual scene. These ratings are pooled together into a smooth map, which is supposed to capture the distribution of meaning across the image. Compared to outputs from a simple saliency model (GBVS, Harel et al., 2006), MMs were more predictive of human fixations. On that basis it has been claimed that meaning guides human fixations in natural scene viewing (Henderson & Hayes, 2017, 2018). Here, we examined central predictions of this claim.

First, if MMs measure meaning and if meaning guides human eye-movements, MMs should be better in predicting locations of fixations than saliency models because these models rely solely on image features. Therefore, we compared MMs to a range of classic and state-of-the-art models. We replicate the finding that MMs perform better than some of the most basic saliency models. Contrary to the prediction, however, DeepGaze II (DGII; Kümmerer, Wallis, & Bethge, 2016; Kümmerer et al., 2017), a model based on a deep convolutional neural network, outperforms MMs.

A second prediction is that if MMs are sensitive to meaning and if meaning guides human gaze, differences in eye movements that result from changes in meaning should be reflected in equivalent differences in MMs. We probed this prediction experimentally using a well-established effect: the same object, when presented in an atypical context (e.g., a shoe on a bathroom sink) attracts more fixations than when presented in a typical context because of the change in the semantic object-context relationship (Henderson, Weeks, & Hollingworth, 1999; Öhlschläger & Võ, 2017). Replicating previous studies, image regions attracted more fixations when they contained context-inconsistent compared to context-consistent objects. Crucially, however, MMs of the modified scenes did not attribute more 'meaning' to these regions. DGII also failed to adjust its predictions accordingly.

Together, these findings suggest that semantic information contained in visual scenes is critical for the control of eye movements. However, this information is captured neither by MMs nor DGII. We suggest that similar to saliency models, MMs index the distribution of visual features rather than meaning.

## Method

We conducted a single experiment in which human observers free-viewed natural scenes while their eye-movements were being recorded. The obtained data was analyzed in two complimentary ways. First, we compared how well MMs and different saliency models predict locations of human fixations in natural scenes. Subsequently, we assessed the sensitivity of MMs and the best-performing saliency model to manipulations of scene meaning. The reported experiment was not preregistered. The data, the code to create MMs, and all openly available resources used in the study can be accessed via the links provided in the Supplement.

### Stimuli

We used images from two conditions of the SCEGRAM database (Öhlschläger & Võ, 2017): the Consistent and the Semantically Inconsistent conditions (called ‘Inconsistent’ here). In the Consistent condition (used in both analyses), scenes contain only objects that are typical for a given context. In the Inconsistent condition (used only in the second analysis), one of the objects is contextually inconsistent. For example, a hairbrush in the context of a bathroom sink from the Consistent condition is replaced with a flip-flop in the Inconsistent condition (see Figs. 1a and 1b). Such changes in object-context relationship alter the meaning attached to the manipulated object. For every scene, we indexed the location of the consistent and inconsistent objects with the superimposed bounding boxes for both objects (see Figs. 1a and 1b). We refer to this location as the Critical Region, because it is the only part of the image that changes between Consistent and Inconsistent conditions. We used 36 selected scenes in both conditions (72 photographs in total, listed in the Supplement together with the selection criteria). We also replicated the main finding of the first analysis in an additional set of 30, very different, images (reported in the Supplement).

**Fig. 1.**
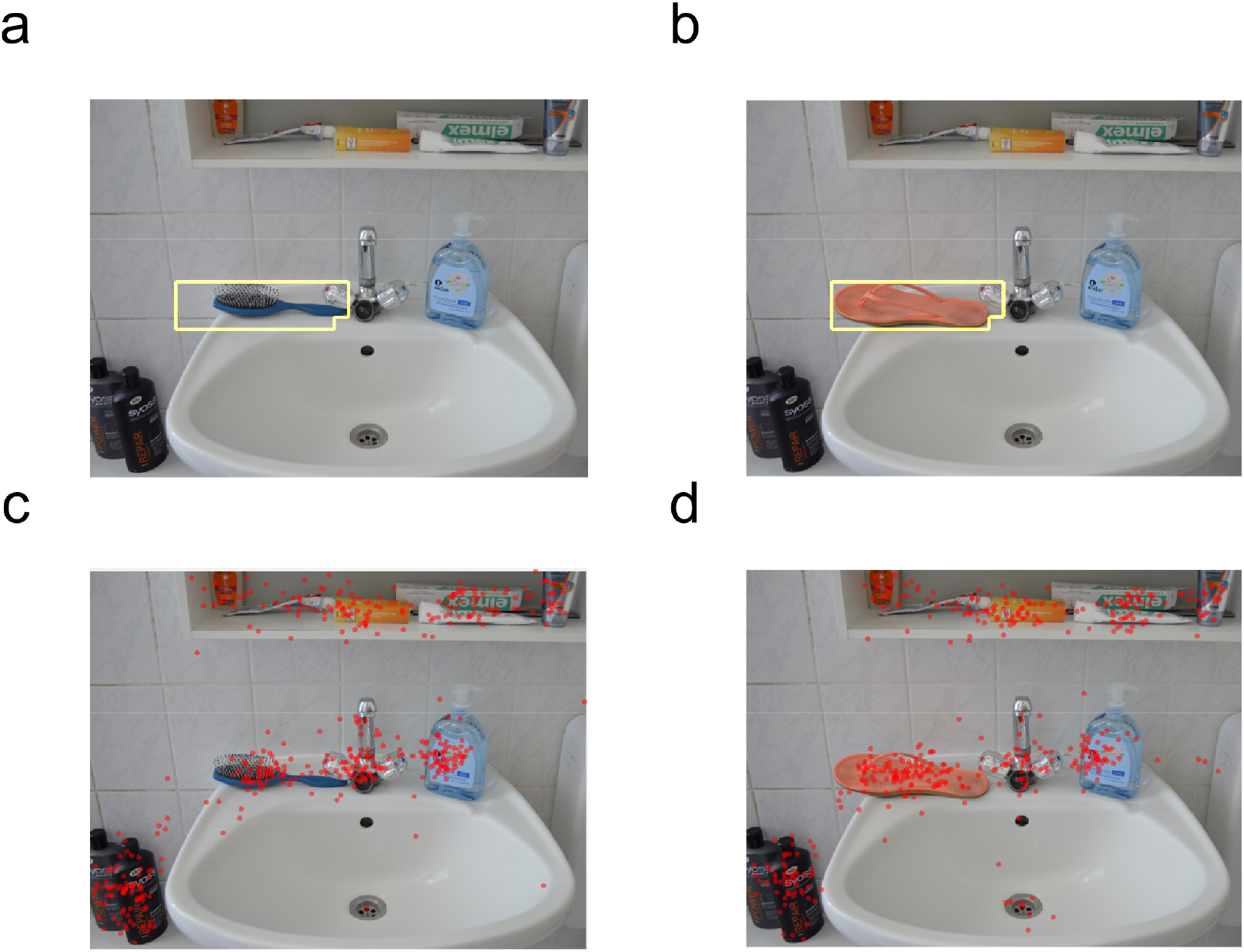
Illustration of sample stimuli in (a) the Consistent and (b) the Inconsistent condition with the Critical Region outlined in yellow and (c, d) human fixations recorded in both conditions. In this example, a hair brush on a bathroom sink (a) – an object consistent with the scene context – has been exchanged for a shoe (b) to introduce semantic inconsistency.

### Procedure

The procedure consisted of 3 blocks, interleaved with breaks. Participants were instructed to ‘look carefully at each’ image. Experimental blocks began with an eye tracker calibration/validation. Within each block, observers free-viewed a series of 24 photographs from both SCEGRAM conditions, each for 7 seconds. After image offset, observers were required to press a button to view the next image. Then, a fixation point appeared centrally on a screen and once observers fixate on it (as determined online by their eye-trace), the actual image was displayed. Before starting the experiment, observers viewed a sample image in an identical regime to familiarize themselves with the procedure. Each stimulus was shown once and the order of presentation was fully randomized. The stimuli were presented against a uniform grey background and had a width of 688 pixels and a height of 524 pixels, which subtended approximately 19.7 and 15 degrees of visual angle, respectively. Stimulus presentation time and size were adopted from a previous study with the SCEGRAM database (Öhlschläger & Võ, 2017).

### Observers

20 volunteers (3 male; mean age 19.4) recruited from the Cardiff University undergraduate population took part in the study. All reported normal or corrected-to-normal vision, provided written consent, and received course credits in return for participation. The study was approved by the Cardiff University School of Psychology Research Ethics Committee. The primary units of interest in our analyses were the distributions of fixations over images. The number of observers we recruited guarantees that including more observers would not change these distributions significantly (demonstrated in the Supplement).

### Apparatus

The study was conducted in a dimly lit room. SCEGRAM images from both conditions were presented on an LCD monitor (Iiyama ProLite B2280HS, resolution 1920 by 1080 pixels, 21 inches diagonal). Chin and forehead rests were used to ensure that observers maintained the constant distance of 49 cm from the screen. Their eye movements were recorded with the frequency of 500 Hz using an EyeLink 1000+ eye tracker placed on a tower mount. The experiment was controlled by custom-written Matlab (R2017a version) scripts using Psychophysics Toolbox Version 3 (Kleiner, Brainard, & Pelli, 2007).

This procedure was conducted separately for the fine and coarse grid, and the meaning map for a given image was created by averaging the two outcomes and normalizing the result to a range between 0 and 1.

### Creating MMs

To create MMs for our stimuli, we followed the procedure described by Henderson & Hayes (2017, 2018; for details see Fig. 2). Each image was segmented into partially overlapping patches of two sizes: fine patches had a diameter of 107 pixels (3 degrees of the visual angle, or 16 % of the image width), coarse patches of 247 pixels (7 degrees or 36% of the image width) (Fig. 2a and b). Their centers were 58 pixels (fine) and 97 pixels (coarse) apart from each other.

**Fig. 2.**
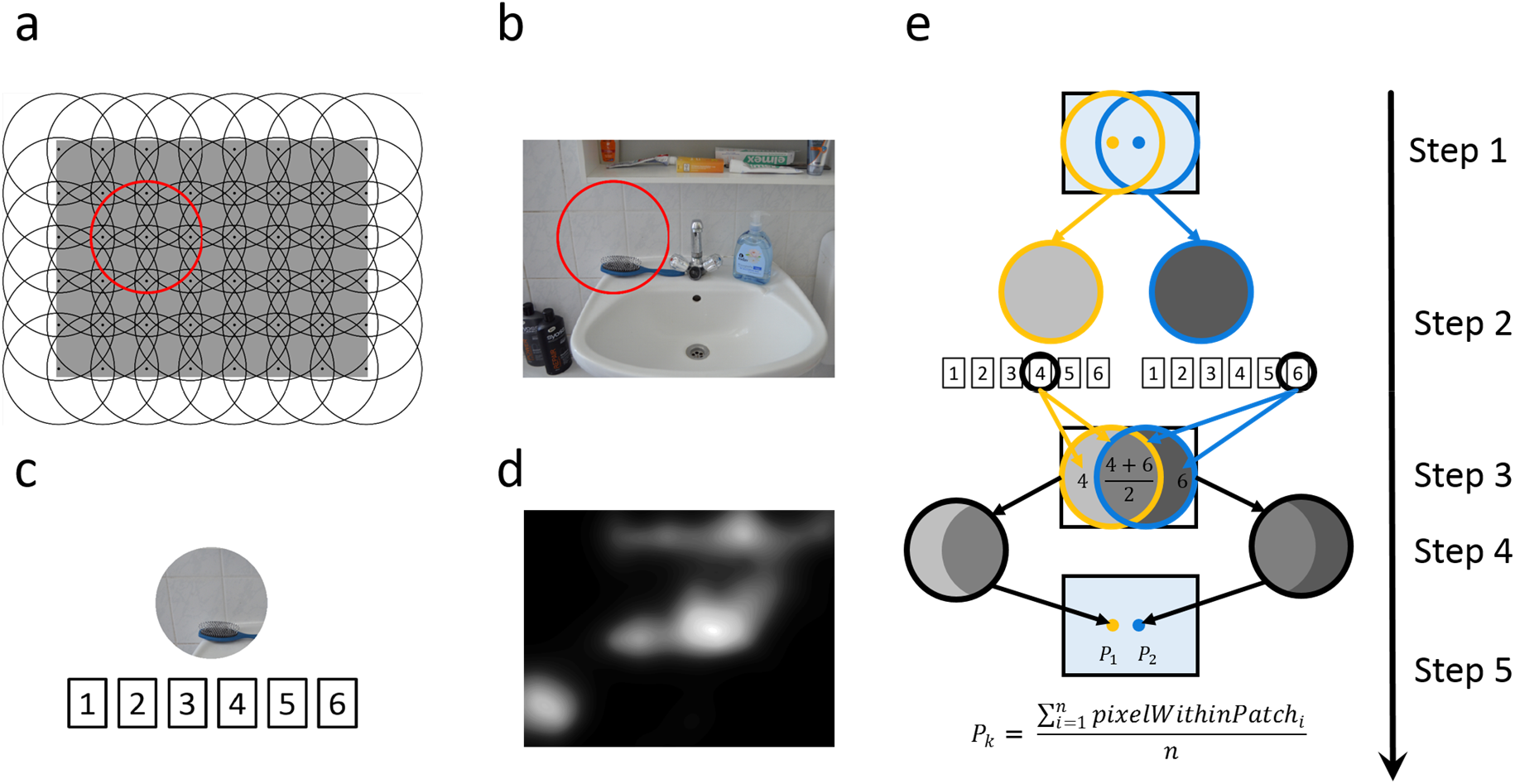
Illustration of the stimuli and procedure used for creating meaning maps. (**a**) Grids of equally spaced circles were used to cut images into fine and coarse patches (only the latter are illustrated here). The red circle indicates a sample patch in the grid. (**b**) Here, the sample patch is highlighted in one of the scenes from the Consistent condition. (**c**) Patches were presented in isolation and rated for their meaningfulness by three independent observers on a scale from 1 to 6. The panel has illustrative purpose only – the scale presented to observers included additional labels (ranging from ‘Very Low’ to ‘Very High’). (**d**) Illustration of a meaning map with greyscale values indicating ‘meaningfulness’. (**e**) Simplifying illustration of how meaning maps are generated from ratings. For simplicity sake, only two patches are shown (step 1). Each patch is rated in isolation (step 2; here only one rating per patch is shown). All pixels within an image area are then assigned average rating values, taking into account all ratings for patches that overlap with this area (step 3). For the area of the original patch (step 4), all pixels are then averaged and the resulting value is assigned to the center of the patch (step 5). Finally, the patch centers were used as interpolation nodes for thin-plate spline interpolation producing a smooth distribution of values over the image (not illustrated).

Next, we collected meaningfulness ratings from human subjects for all patches. Each patch was presented in isolation and rated for its meaningfulness on a 6 point Likert scale (Fig. 2). As in Henderson and Hayes (2017), we used a Qualtrics survey completed by naive observers recruited via the crowdsourcing platform Amazon Mechanical Turk (see Supplement for eligibility criteria). Each participant provided ratings for 305 or 303 patches of both sizes (selected randomly from all images), on average spent approximately 14 min on the task, and received 2.18 USD as remuneration. In total, 69 individuals were used as raters, with three individuals rating each individual patch. The collected ratings were then used to create MMs (see Fig. 2).

When creating MMs for images from both conditions, we exploited the fact that photographs from the Consistent and Inconsistent conditions differ only in the Critical Region (the part of the image containing the manipulated object) while the remaining parts overlap. We collected meaningfulness ratings for the patches belonging to overlapping areas only once, and the separate sets of ratings for Consistent and Inconsistent condition were collected only for those patches that contained at least one pixel belonging to the Critical Region. In total, the number of patches rated in the study amounted to 7013: 4840 fine patches (of which 520 belonged to the images from the Inconsistent condition) and 2173 coarse patches (445 Inconsistent).

### Saliency models

In the first analysis, we compared predictive performance of MMs to four saliency models of different complexity. The first two models – GBVS (Harel et al., 2006) and AWS (Garcia-Diaz, Fdez-Vidal, Pardo, & Dosil, 2012) – rely on simple visual features, such as local colors and edge orientations, and share the assumption that fixations land on image regions distinct from their surroundings in terms of values of these features. By contrast to GBVS, AWS includes a statistical whitening procedure to improve performance. Both these models were previously used to estimate the influence of image features relative to cognitive factors on the deployment of fixations: GBVS in the previous studies with MMs, AWS elsewhere (Stoll et al., 2015).

Two other models that we compared to MMs – ICF and DeepGaze II (DGII) – were designed in a data-driven manner (Kümmerer et al., 2017). Both have the same architecture, consisting of a fixed network that extracts sets of features from images and a readout network that is trained on human fixations separately for each model to combine the features in a way to maximize the models’ predictive power. While the fixed network of ICF extracts only simple visual features (local intensity and contrast), DGII is tuned to features extracted by a deep convolutional neural network pre-trained for object recognition (VGG-19; Simonyan & Zisserman, 2014).

All saliency models output smooth maps that predict the probability of image regions to be fixated. Human observers have the tendency to look at the center of images (Tatler, 2007), and therefore this probability is usually higher in the central region of the image. This ‘center bias’ has important consequences for the evaluation of saliency models. Their performance differs depending on whether they are evaluated using a metric expecting some form of this bias or not (Kümmerer, Wallis, & Bethge, 2018). Here, for the sake of simplicity, we do not incorporate center bias in the models or in the MMs (unlike the original authors) and use an appropriate metric for this situation (see Performance metrics section). Importantly, analyses addressing the issue of center bias in a more extensive way (reported in the Supplement) provide only further support for our conclusions.

### Data pre-processing

Fixation locations from the eye tracker recordings were extracted using the algorithm provided by the device manufacturer operating with the default parameter values. Thereby, we obtained a discrete distribution of fixations on each image (see Fig. 1c and 1d). Then, in line with the previous MMs studies, we smoothed these discrete distributions with a Gaussian filter with a cutoff frequency of −6 dB, using the function provided by Bylinskii and colleagues (2014).

Next, smooth distributions from fixations, models, and MMs were separately normalized to a range from 0 to 1 for each image. Finally, for each scene, histograms of all distributions from both conditions were matched to histograms of smoothed fixations from Consistent condition using the Matlab imhistmatch function, as in the original MMs studies. Histogram matching makes distributions directly comparable as it ensures that they differ only with respect to their shape, and not their total mass.

### Performance metrics

To compare the ability of MMs and models to predict locations of human fixations in Experiment 1, we use two well-established metrics (Bylinskii, Judd, Oliva, Torralba, & Durand, 2016): Correlation and Shuffled Area Under ROC curve (sAUC; Zhang, Marks, Tong, Shan, & Cottrell, 2007) with the implementations provided by Bylinskii and colleagues (2014).

Correlation, used in the previous studies on MMs, is calculated as Pearson's linear correlation coefficient between a smoothed distribution of observers’ fixations over the image and predictions of a saliency model or MMs. We additionally used sAUC (Zhang et al., 2008), which, unlike Correlation, guarantees that the measured differences in performance between models are driven by their sensitivity to factors guiding fixations, and not by the degree to which they include human center bias in their predictions, even implicitly (Kümmerer, Wallis, & Bethge, 2015; Kümmerer et al., 2018).

## Comparing meaning maps and saliency models – results

In the first analysis, we compared performance of four saliency models to MMs in predicting human fixations in the Consistent condition, i.e., when viewing typical scenes with no obvious object-context inconsistencies (Tab. 1, Fig. 3). If human gaze is guided by meaning, and if MMs provide an index for the distribution of meaning, we would expect MMs to outperform all saliency models because these models are based solely on image features.

**Table 1.**
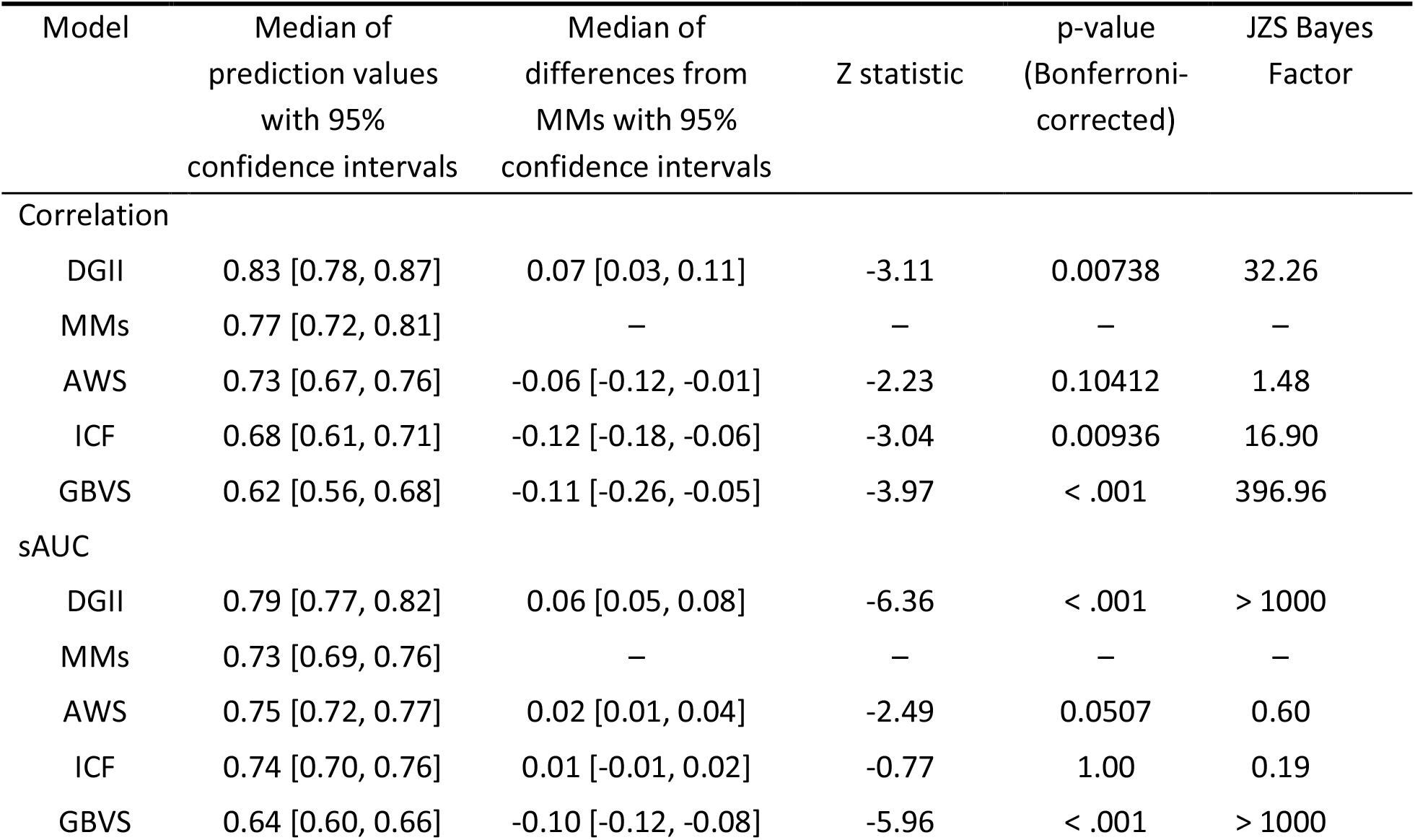
Comparison of Predictive Power of Saliency Models and MMs Using Correlation and sAUC.

**Fig. 3.**
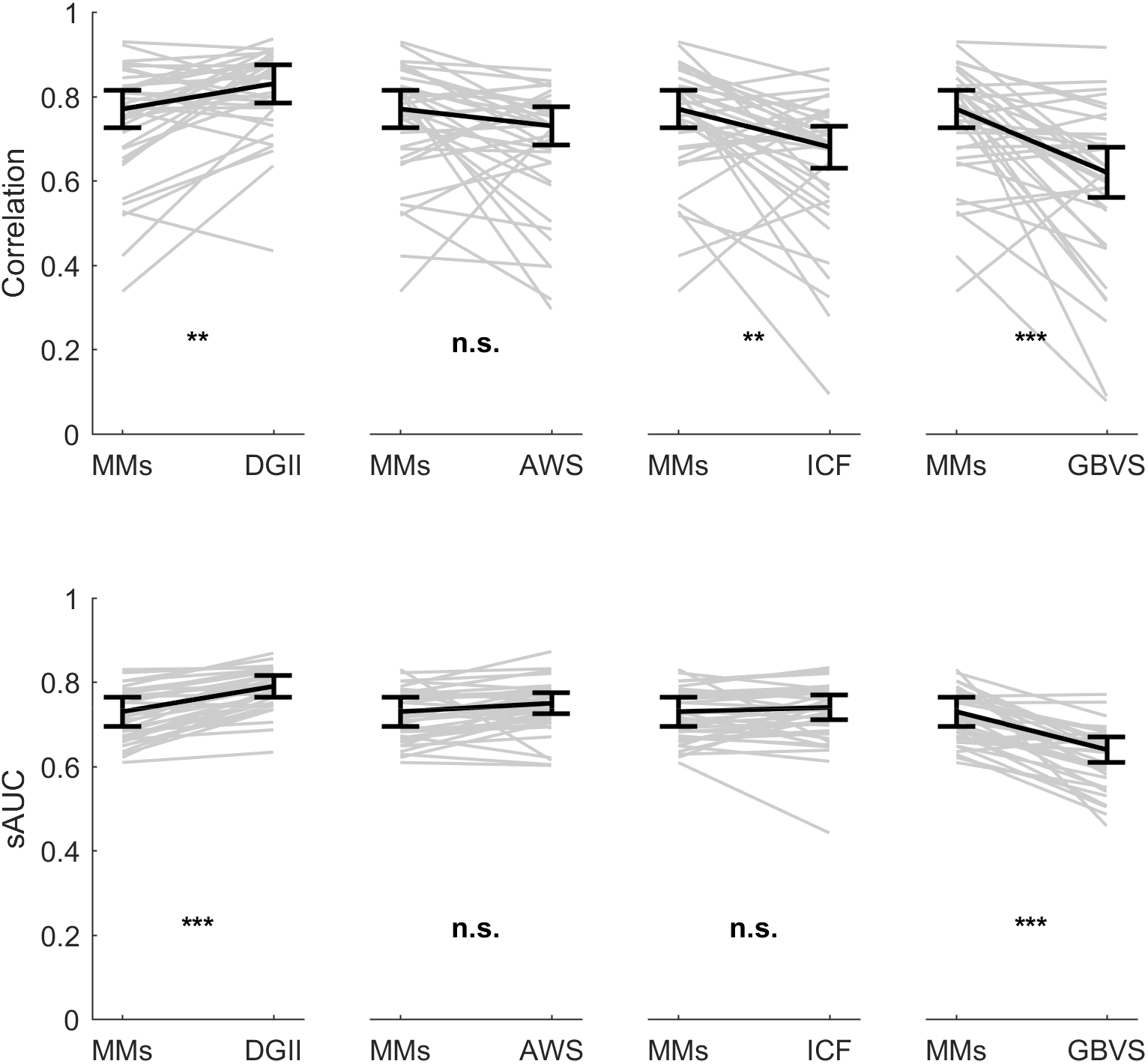
Performance of MMs and saliency models in predicting human fixations according to (a) Correlation and (b) sAUC metrics. Note that according to both metrics DGII predicted human fixations better than MMs. Asterisks indicate p-values from statistical tests comparing MMs to different models (reported in Table 1.): * indicates p ≤.05, ** p ≤.01, *** ≤.001 and ‘n.s.’ indicates the lack of statistical significance. Grey lines connect values obtained for individual images. Black vertical bars indicate 95% confidence intervals for the medians

### Predictive power

Correlation and sAUC values obtained for MMs and for each of the models were compared using Bonferroni-corrected paired Wilcoxon tests (Fig. 3; Tab. 1). We used non-parametric tests because for some of the distributions the assumptions of normality was not met. For the same reason we chose a median as a measure of centrality (we calculate confidence intervals for median using a bootstrapping method – see details in the Supplement). Additionally, we calculated JZS Bayes Factor (Rouder, Speckman, Sun, Morey, & Iverson, 2009) to quantify the evidence for (or against) the differences between models and MMs (Tab. 1). While deviations from normality can be problematic for Bayes factor analyses, they are most likely not an issue in the current situation: the Bayes factors for the key finding are large and the deviations from normality are small.

As shown in Tab. 1 and on Fig. 3, according to both measures, MMs outperformed GBVS in predicting human fixations, thereby replicating the results of Henderson and Hayes (2017, 2018) using new images and new participants. Contrary to expectations, however, both metrics indicated that DGII predicted fixations better than MMs. Furthermore, performance of AWS and MMs did not differ significantly irrespective of the metrics. Finally, MMs outperformed ICF according to Correlation, but not sAUC. In fact, for the latter metric, JZS-Bayes Factor indicated support for the null hypothesis.

### Semi-partial correlations

Because predictions of models and MMs overlap, we quantified their distinct predictive power using semi-partial correlations. We conducted these analyses for GBVS (used in the original MMs studies) and DGII (the only model which markedly outperformed MMs).

For each scene from the Consistent condition, we calculated two semi-partial correlations with the distribution from smoothed fixations: one for MMs while controlling for GBVS, and one for GBVS while controlling for MMs (see Fig. 4). Consistent with findings by Henderson and Hayes (2018), MMs explain more unique variance than GBVS (Fig. 6a), as indicated by the significantly higher coefficients in the former than the latter case (mean difference 0.28, 95% confidence interval (CI) [0.17, 0.39]; paired t-test, t(35) = 5.22, p < .001). Interestingly, the identical analysis with DGII revealed that DGII explained significantly more unique variance than MMs (mean difference 0.15, 95% CI [0.07, 0.24]; t(35) = 3.60, p < .001, see also Fig. 4b).

**Fig. 4.**
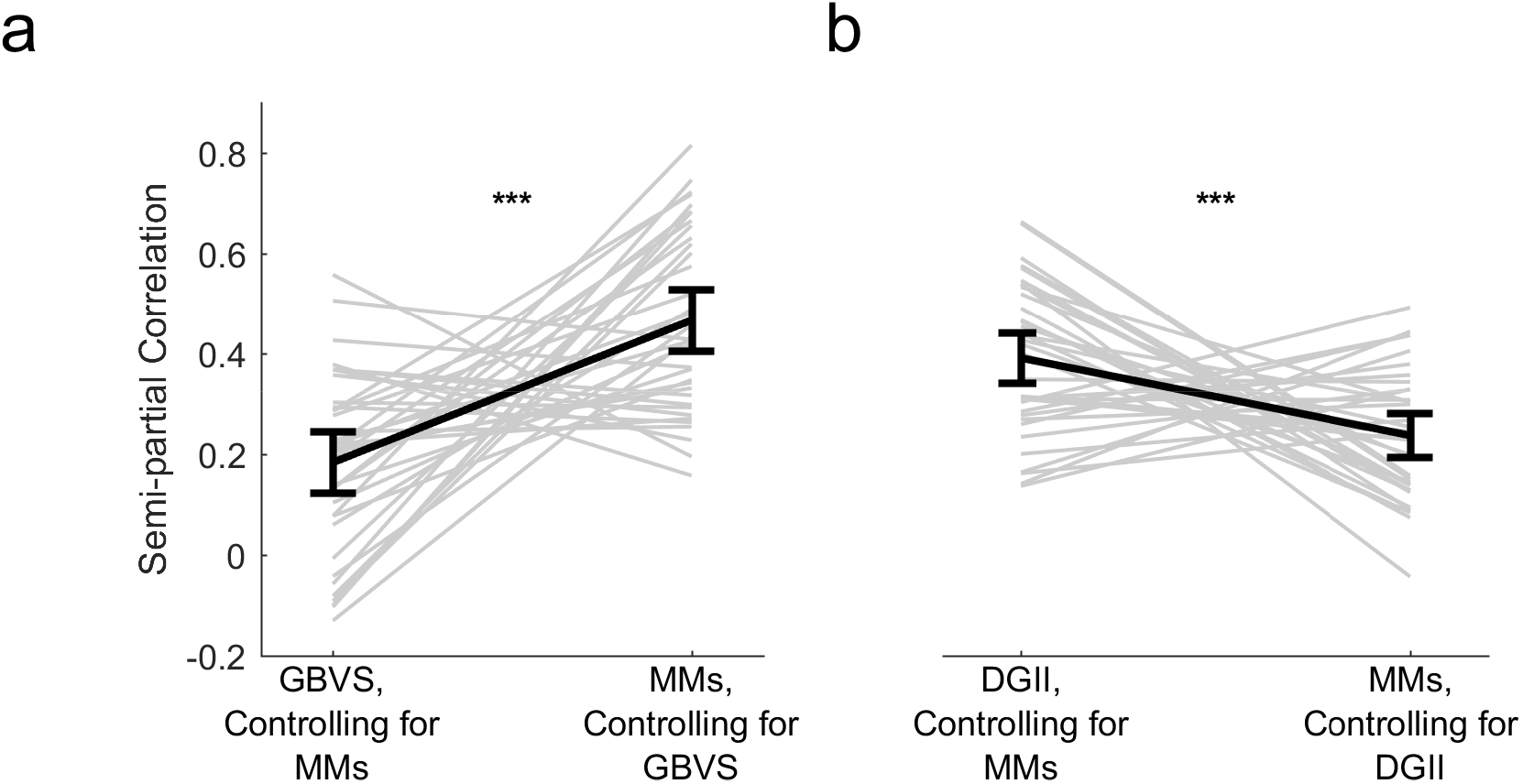
Comparison of semi-partial correlations with smoothed human fixations for (a) MMs and GBVS and for (b) MMs and DGII. The obtained coefficients were significantly higher when assessing MMs while controlling for GBVS compared to when assessing GBVS when controlling for MMs. The opposite was true for the analyses with DGII. All figure characteristics are as in Fig. 3. except that medians instead of means are presented.

### Internal replication

To demonstrate the generalizability of our conclusions beyond SCEGRAM images, we replicated the main results with a different stimulus set (see the Supplement).

## Comparing meaning maps and saliency models – discussion

If human gaze is guided by meaning, and if MMs index the distribution of meaning across an image, MMs should outperform saliency models that are exclusively based on image features. Our first analysis showed that this prediction does not hold. In fact, DGII generated better predictions and explained more unique variance than MMs. Therefore, at least one of the two premises of our prediction is wrong: either human eye-movements are not sensitive to meaning or MM do not index meaning. The second analysis allowed us to distinguish between these alternatives.

## Analyzing the effects of semantic inconsistencies within scenes – method

In the second analysis, we assessed how human observers, DGII, and MMs respond to experimental changes in meaning induced by altered object-context relationships. We used eye-movement data from both the Consistent and the Inconsistent condition. These conditions differed solely in the Critical Region, an area that either contained an object that was either consistent with the scene context or induce semantic conflict. For each scene, we calculated the mass of the distributions of human gaze, DGII, and MMs falling into the Critical Region, respectively, and divided it by the Region’s area for normalization. Our primary interest was the comparison between conditions: to the extent to which humans, DGII, and MMs are sensitive to meaning, they should fixate more (humans) or predict more fixations (DGII and MMs) on the Critical Region in the Inconsistent than the Consistent condition.

## Analyzing the effects of semantic inconsistencies within scenes – results

Our comparison indicated that, as predicted, observers fixated more on inconsistent than consistent objects (Fig. 5a). By contrast, behavior of both MMs and DGII did not change across conditions (Fig. 5b and c). These impressions were confirmed by a 2×3 ANOVA, with condition (Consistent vs. Inconsistent) as a within-subjects factor and the distribution source (human fixations vs. MMs vs. DGII) as a between-subjects factor. We found a statistically significant main effect of distribution source, F(2, 105) = 13.09, p < .001, ω^2^= 0.16 and condition, F(1, 105) = 7.41 p = 0.0076 X, ω^2^ = 0.005. These main effects were qualified by a significant interaction, F(2, 105) = 16.90, p < .001 X, ω^2^ = 0.026. Tukey post-hoc tests showed that human observers looked more at the Critical Regions in the Inconsistent, than the Consistent condition, t(105) = −6.22, p < .001. In contrast, no significant differences between conditions were found for DGII, t(105) = −0.09 p = 1.0, and MMs, t(105) = 1.60 p = 0.6028. Comparisons within conditions indicated that human fixations differed from MMs in the Inconsistent condition, t(129.91) = 5.78 p < .001, but not the Consistent condition, t(129.91) = 2.16 p = 0.2662. A significant difference between DGII and human fixations was detected in both Consistent, t(129.91) = −2.96 p = 0.0420, and Inconsistent conditions, t(129.91) = −5.79 p < .001.

**Fig. 5.**
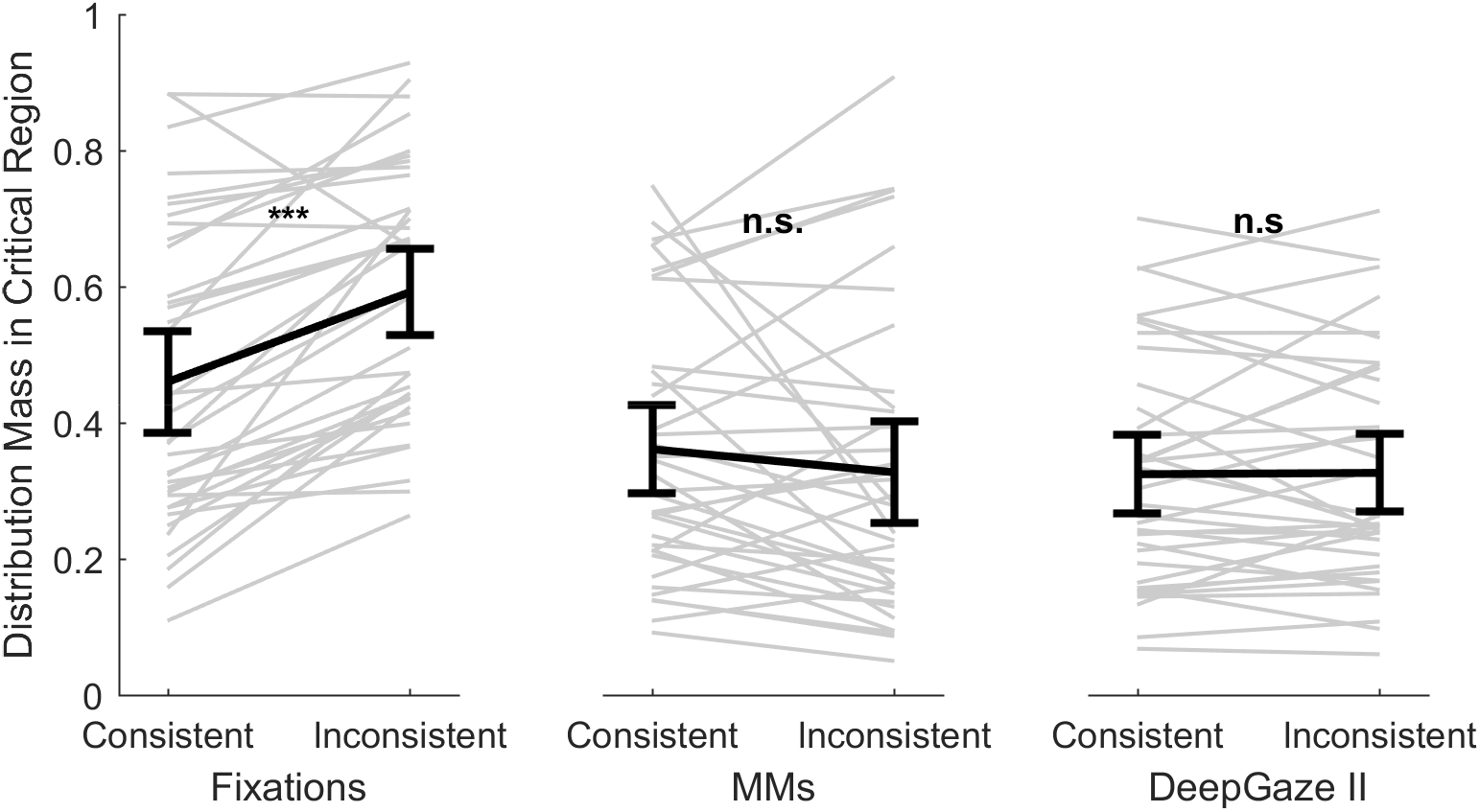
Normalized distribution mass falling within Critical Regions in both conditions for (a) smoothed human fixations, (b) MMs, and (c) DGII. All figure characteristics are as in Fig. 3.

Additionally, conditions differed regarding the number of fixations per image, t(35) = 5.67 p < .001. On average, there were 6% fewer fixations in the Inconsistent condition. This excludes the possibility that higher number of fixations in this condition might drive the observed increase in the distribution mass falling within the Critical Regions.

Finally, systematic differences in object size between Consistent and Inconsistent conditions could affect our results because larger objects may attract more fixations solely because they occupy a larger image area. However, this factor was minimized by showing each object in a consistent and an inconsistent context. Yet, the same object might be shown in a slightly different position in the two conditions and might therefore occupy slightly different amounts of the image. This was, however, not the case: the JZS Bayes Factor of 4.26 indicated that the two conditions did not differ in the size of the bounding boxes of each manipulated object (objects in the Inconsistent condition were on average 1562.28 pixels larger; 95% confidence interval: [−2582.74, 5707.29]).

To summarize, semantic changes induced by altering object-context relationships elicited changes in distributions of human fixations, but neither MMs nor DGII could predict them. These results suggest that both models might be sensitive to image features, which are frequently correlated with image meaning, rather than to meaning itself.

## Discussion

A long-standing debate in visual perception concerns the extent to which visual features vs. semantic content guide human eye-movements in free viewing of natural scenes. To distinguish these hypotheses, indexing the distributions both of features and meaning across an image is critical. While image-based saliency models have been used to index features for two decades, measuring semantic importance has been difficult until meaning maps (MMs) have recently been proposed. Here, we assessed the extent to which MMs indeed capture the distribution of meaning across an image. First, we demonstrate that despite the purported importance of meaning as measured by MMs for gaze control, MMs are not better predictors of locations of human fixations than at least some saliency models, which are based solely on image features. In fact, DeepGaze II (DGII), a model using deep neural network features, outperformed MMs. Second, we assessed the sensitivity of human eye-movements, MMs, and DGII to changes in image meaning induced by violations of typical object-context relationships. Observers fixated more often on regions containing objects inconsistent with scene context (thus replicating previous findings) but these regions were not indexed as more meaningful by MMs, or as more salient by DGII. Together, these findings challenge central assumptions of MMs, suggesting that they are insensitive to the semantic information contained in the stimulus.

The good performance of DGII in predicting human gaze might be attributable to the high-level features it extracts from images. Three other models, which use low-level features, failed to decisively outperform MMs. However, unlike two of them (GBVS and AWS), DGII is trained with data on human fixations to optimize performance (Kümmerer et al., 2016, 2017). Yet, training alone cannot explain the difference in performance. The third low-level feature model (ICF) is trained in the same way (Kümmerer et al., 2017) but still achieves a lower performance than DGII. These findings suggest that feature type is indeed critical for a model’s performance. Importantly, however, while DGII uses high-level features transferred from a deep neural network trained on object recognition (Simonyan & Zisserman, 2014), this is not equivalent to indexing meaning. Rather, the good performance of DGII is likely due to meaning supervening on, or correlating with, some of the features indexed by this model.

Correlation between visual features and meaning as the source of good performance in saliency models has already been considered by the authors of MMs (Henderson & Hayes, 2017). Our findings suggest that MMs might share this characteristic with saliency models. Specifically, the ratings used to construct MMs might be based on visual properties in such a way that highly structured patches that contain high-level features receive high ratings. These features often correlate with meaning, but in and of themselves do not amount to meaning. According to this interpretation, both DGII and MMs index high-level features. Their success in predicting human behavior derives from the typically strong correlation between high-level features and meaning, with a higher correlation for the features extracted by DGII than MMs.

An alternative interpretation of the finding that DGII outperforms MMs is that image features rather than meaning guide human fixations. However, this interpretation is inconsistent with our second analysis. Here, observers clearly exhibited sensitivity to meaning, as indicated by changed gaze patterns after introducing semantic inconsistencies into the scenes. This experimental manipulation targets a type of meaning that is based on how objects relate to the broader context in which they occur. While specific, it is precisely this kind of meaning that is of high theoretical importance in eye-movement research (Henderson, 2017; Henderson et al., 2009). Thus, even if MMs were to measure other types of meaning, as has been suggested (Henderson et al., 2018), the fact that they are not sensitive to meaning derived from object-context relationships seriously limits their usefulness. Moreover, the idea that MMs indeed index other kinds of meaning that are important for guidance of fixations is not consistent with our findings. If this were the case, then we would expect MMs to predict human fixations better than saliency models that solely rely on image features, which is not the case.

The insensitivity to semantic inconsistencies reveals inherent limitations of both MMs and DGII. The way in which MMs are constructed implicitly assumes that meaning is a local image-property, which is not true for object-context (in)consistency. This limitation may potentially be alleviated by ‘contextualized MMs’ (Peacock, Hayes, & Henderson, 2019), a recently suggested modification of the ‘standard’ MMs. These novel maps are created from meaningfulness ratings by observers who see the whole scenes from which the to-be-rated patches were derived. It is yet to be seen what this approach can reveal about fixation selection beyond the fact that humans asked to indicate meaningful or interesting regions within scenes highlight areas, which tend to be frequently fixated by other observers (Nyström & Holmqvist, 2008; Onat et al., 2014). DGII, in turn, does not explicitly encode semantic information, and was not trained on the relationship between eye movements and semantic (in)consistency. But its failure highlights an opportunity to improve saliency models by incorporating semantic relationships (Bayat, Koh, Nand, Pereira, & Pomplun, 2018).

Taken together, our results suggest that, contrary to their core promise as a methodology, meaning maps (MMs) do not offer a way to measure the spatial distribution of meaning across an image. Instead of meaning per-se, they seem to index high-level features that have the potential to carry meaning in typical natural scenes. They share this characteristic with state-of-the-art saliency models, which are easier to use, do not require human annotation, and yet predict locations of human fixations better than MMs.

## Author contributions

C.T. and M.P. conceived of the study. M.P., T.W., and C.T. designed the experiment. M.P. collected and analyzed the data under the supervision of C.T. with support from M.K., T.W., and M.B. The paper was drafted by M.P. and C.T.; T.W., M.K., and M.B. provided detailed comments.

