## Supplement for "Meaning maps and saliency models based on deep convolutional neural networks are insensitive to image meaning when predicting human fixations"

#### Internal replication

We replicated the finding that DGII outperforms MMs in predicting human fixations in a separate experiment with a different set of stimuli and new observers. This second experiment was identical to the first one except for the stimuli, the number of observers (21 instead of 20), and the number and the duration of images presented.

**Methods.** We used 30 photographs from Corel Photo Library depicting mainly single animals. The images were converted to grayscale and resized to 788 by 526 pixels (22.5 by 15 degrees of visual angle). We presented them for 3 seconds each, in blocks of 10. 21 new observers took part in the study (1 male, mean age 19.05). To create MMs, the images were fragmented into 4200 fine and 1620 coarse patches in total. Each rater recruited via Amazon Mechanical Turk rated 291 patches. Because DGII requires RGB images as input, to generate model's predictions we converted the stimuli to RGB by copying greyscale pixel values to the three color channels.

**Results.** DGII again outperformed MMs in predicting human fixations (see Table S1).

Table S1. Comparison of Predictive Power of Saliency Models and MMs Using Correlation and sAUC – Internal Replication

| Model | Median of prediction values with 95% CIs | Median of differences from MMs with 95% CIs | Z statistic | p-value (Bonferroni-corrected) | JZS Bayes Factor |
|---|---|---|---|---|---|
| Correlation | | | | | |
| DGII | 0.86 [0.71, 0.89] | 0.11 [0.04, 0.17] | -3.28 | 0.0042 | 6.34 |
| MMs | 0.74 [0.62, 0.80] | – | – | – | – |

| | | | | | |
|---|---|---|---|---|---|
| sAUC | | | | | |
| DGII | 0.75 [0.73, 0.78] | 0.04 [0.02, 0.06] | -3.87 | < .001 | 63.52 |
| MMs | 0.70 [0.68, 0.74] | – | – | – | – |

**Analysis including optimal center bias**

Comparison between saliency models is complicated by the fact that different performance metrics can result in different rankings (Bylinskii et al., 2014; Kümmerer, Wallis, & Bethge, 2018). This problem can be mitigated by a post-processing procedure that optimizes model predictions for assessment with a specific metric (Kümmerer, Wallis, & Bethge, 2015; Kümmerer et al., 2018). This procedure estimates the optimal monotone nonlinearity to be applied to prediction value of each pixel and the optimal center bias to be added to model predictions. Parameter optimization proceeds according to the maximum-likelihood principle. This approach levels the differences between metrics and ensures that a central bias is modelled optimally for each metric (Kümmerer et al., 2018).

We post-processed MMs and predictions of the saliency models in this way. Since our dataset is smaller than the one used by Kümmerer and colleagues (2015), we used a smaller number of parameters (8 for the nonlinearity and 6 for the center bias). To avoid overfitting, predictions for each image were processed (separately for Correlation and sAUC) with parameters optimized on predictions for all other images. According to both metrics, including post-processing did not change the main result – DGII still outperformed MMs in predicting human fixations (see Table S2.).

Table S2. Comparison of Predictive Power of Saliency Models and MMs Using Correlation and sAUC – Analysis Including Optimal Centre Bias

| Model | Median of prediction values with 95% | Median of differences from MMs with 95% | Z statistic | p-value (Bonferroni-corrected) | JZS Bayes Factor |
|---|---|---|---|---|---|

| | confidence intervals | confidence intervals | | | |
|---|---|---|---|---|---|
| Correlation | | | | | |
| DGII | 0.87 [0.81, 0.89] | 0.08 [0.04, 0.15] | -4.68 | < .001 | > 1000 |
| MMs | 0.77 [0.72, 0.80] | – | – | – | – |
| AWS | 0.79 [0.73, 0.82] | 0.02 [-0.02, 0.04] | -0.63 | 1 | 0.18 |
| ICF | 0.75 [0.69, 0.77] | - 0.02 [-0.09, 0.01] | -1.05 | 1 | 0.34 |
| GBVS | 0.62 [0.59, 0.69] | - 0.11 [-0.20, -0.04] | -4.27 | < .001 | > 1000 |
| sAUC | | | | | |
| DGII | 0.81 [0.78, 0.83] | 0.05 [0.03, 0.06] | -6.55 | < .001 | > 1000 |
| MMs | 0.77 [0.73, 0.78] | – | – | – | – |
| AWS | 0.77 [0.74, 0.78] | 0.01 [0.00 0.03] | -1.37 | 0.6850 | 0.36 |
| ICF | 0.71 [0.69, 0.72] | -0.05 [-0.06 -0.04] | -5.03 | < .001 | > 1000 |
| GBVS | 0.65 [0.61, 0.67] | -0.11 [-0.13 -0.09] | -6.55 | < .001 | > 1000 |

**Number of observers**

We tested if fixations from 20 observers who viewed images in our experiment are sufficient to closely approximate the theoretical ground truth distributions of fixations which would have been obtained from an infinite number of observers. Visual inspection of the data revealed that including fixations from about 10 observers results in distributions which remain virtually unchanged when fixations from more observers are added. This observation was confirmed by a more formal analysis. For each observer, we randomly selected a subset of 12 other observers 10 times and calculated how well smoothed fixations from these subsets predict fixations of the observer for each image using correlation. Averaging all the obtained values over observers and subsets resulted in an estimate of how well 12 observers predict fixations of a single observer viewing a given image. The value obtained for all SCEGRAM images amounted to 0.835 (SD = 0.094). Next, again for each observer viewing each image, we calculated how well their fixations can be predicted using fixations of the remaining 19 observers. The average of the obtained values equaled to 0.840 (SD = 0.097). This number is close to the value obtained for 12 observers, thus indicating that the underlying distributions of fixations are similar, and that increasing the number of observers would not affect them substantially.

### SCEGRAM scenes

We used SCEGRAM photographs from two conditions: Consistent and (Semantically) Inconsistent. These photographs are divided into quadruples: object A in scene A (scene and object are consistent), object A in scene B (scene and objects are inconsistent), object B in scene B, and object B in scene A. In order to avoid potential distortions of the results, in our experiment we included only these quadruples in which both manipulated objects do not contain any digits or text observers might be trying to read. Applying this selection criterion resulted in retaining the following SCEGRAM scenes: 1-4, 7, 8, 11, 12, 17, 18, 21, 22, 27, 28, 33-44, 47-50, 55, 56, 59-62.

### Eligibility criteria for Amazon Mechanical Turk raters

Raters who rated the meaningfulness of image patches had to meet the following requirements: they had to have an approval rating grater than 96%, be located in the United States, have more than 500 tasks (‘HITs’) approved, and not have completed the task for images from a given set of images (SCEGRAM or from the replication) before.

### Statistical software

All statistical analyses were conducted in R (R Core Team, 2016) with the help of functions from the following packages: BayesFactor (Morey & Rouder, 2015), jmv (The jamovi project, 2019), ppcor (Kim, 2015), and boot (Canty & Ripley, 2019; Davison & Hinkley, 1997).

### Confidence intervals for medians

All confidence intervals for medians were calculated using adjusted bootstrap percentile (BCa) method with 10000 bootstrap replicates, implemented in R package boot (Canty & Ripley, 2019; Davison & Hinkley, 1997).

**Openly available materials:**

Deep Gaze II and ICF: https://deepgaze.bethgelab.org/

AWS: http://persoal.citius.usc.es/xose.vidal/research/aws/AWSmodel.html

GBVS: http://www.vision.caltech.edu/~harel/share/gbvs.php

SCEGRAM database: https://www.scenegrammarlab.com/research/scegram-database/

Code for creating MMs used in this paper: DOI: 10.5281/zenodo.3490592 [to be released]

Data from this study: DOI: 10.5281/zenodo.3490434 [to be released]
